# Electrophysiological signatures of spatial boundaries in the human subiculum

**DOI:** 10.1101/218040

**Authors:** Sang Ah Lee, Jonathan F. Miller, Andrew J. Watrous, Michael Sperling, Ashwini Sharan, Gregory A. Worrell, Brent M Berry, Barbara C. Jobst, Kathryn A. Davis, Robert E. Gross, Bradley Lega, Sameer Sheth, Sandhitsu R. Das, Joel M. Stein, Richard Gorniak, Daniel S. Rizzuto, Joshua Jacobs

**Author notes:** Correspondence: Sang Ah Lee Department of Bio and Brain Engineering, Korea Advanced Institute of Science and Technology 291 Daehak-ro, Yuseong-gu, Daejeon, 34141. Republic of Korea, (e-mail). Joshua Jacobs Department of Biomedical Engineering, Columbia University 1210 Amsterdam Avenue New York, NY 10027, (e-mail).

## Abstract

Environmental boundaries play a crucial role in spatial navigation and memory across a wide range of distantly-related species. In rodents, boundary representations have been identified at the single-cell level in the subiculum and entorhinal cortex of the hippocampal formation. While studies of hippocampal function and spatial behavior suggest that similar representations might exist in humans, boundary-related neural activity has not been identified electrophysiologically in humans until now. Here we present direct intracranial recordings from the hippocampal formation of surgical epilepsy patients while they performed a virtual spatial navigation task. Our results suggest that encoding target locations near boundaries elicited stronger theta oscillations than for target locations near the center of the environment and that this difference cannot be explained by variables such as trial length, speed, or movement. These findings provide the first direct evidence of boundary-dependent neural activity localized in humans to the subiculum, the homologue of the hippocampal subregion in which most rodent boundary cells are found.

**Significance Statement:** Spatial computations using environmental boundaries are an integral part of the brain’s spatial mapping system. In rodents, border/boundary cells in the subiculum and entorhinal cortex reveal boundary coding at the single-neuron level. Although there is good reason to believe that such representations also exist in humans, the evidence has thus far been limited to fMRI studies that broadly implicate the hippocampus in boundary-based navigation. By combining intracranial recordings with high-resolution imaging of hippocampal subregions we identified, for the first time in humans, a neural marker of boundary representation in the subiculum.

## Introduction

Research across a wide range of disciplines has converged on the notion that environmental boundaries strongly influence spatial memory and cognition (Lee, 2017). When animals lose track of where they are, they rely heavily on boundary structures to find their way back to the goal (for review, see Cheng & Newcombe, 2005; Lee & Spelke, 2010; Tommasi et al., 2012). Non-boundary features such as objects and surface properties also influence navigation but are used primarily as beacons (e.g., Lee et al., 2006), contextual cues (e.g., Julian et al., 2015), and error-correcting landmarks in path integration (e.g., Etienne et al., 1996). To explain the effect of boundaries in behavior, theorists have proposed that the 3D structure of the environment provides a reliable basis for metric distance computations in spatial mapping(Cheng, 1986; Gallistel, 1990).

Electrophysiological recordings in the rodent hippocampal formation have shown that the spatial coding by place cells and grid cells is highly influenced by environmental boundaries (O’Keefe & Burgess, 1996; Krupic et al., 2015; Stensola et al., 2015; Lever et al., 2002; Hardcastle et al., 2015). Boundary-based models of place mapping (Hartley et al., 2000; Barry et al., 2006) explain the firing fields of place cells as a sum of distance inputs from nearby boundaries, and the existence of boundary cells in the rodent subiculum (Lever et al., 2009) and border cells in the entorhinal cortex (EC) (Solstad et al., 2008) provide evidence of boundary representations at the single neuron level. Boundary cells are theta-modulated, like other spatial cells, and are characterized by their increased firing in response to nearby boundary structures, such as walls, drop-offs, and traversable gaps on the floor (Lever et al., 2009; Stewart et al., 2014). They develop in rat pups at the same time as place cells and earlier than grid cells (Bjerknes et al., 2014).

Although boundary cells have yet to be found in the human brain, behavioral experiments suggest that we share similar boundary-based navigational mechanisms with other animals. For an extended period in human development, boundaries exert a dominant influence on spatial mapping (Hermer-Vazquez et al., 2001). Such boundaries are not limited to large walls but also include subtle 3D structures such as traversable ridges and curbs (Lee & Spelke, 2008, 2011), similar to the characteristics of boundary cells in rodents discussed above. The use of environmental boundaries can also be seen in adults (Hermer-Vazquez et al., 1999; Hartley et al., 2004), and functional neuroimaging studies have established that boundary-based navigation or imagery engages the hippocampus (Doeller et al., 2008; Bird et al., 2010). Other studies have identified boundary representation of visual scenes and its role in navigation upstream from the hippocampus (Park et al., 2011; Ferrara & Park, 2016; Julian et al., 2016).

Challenges to single-cell recording in humans can be partially bypassed by looking for signatures of neural activity that would be visible at the population level. An example of this is the hexagonally clustered fMRI response in the entorhinal cortex that might be attributed to populations of grid cells (Doeller et al., 2010). Similarly, neural signals of boundary representations could also be visible at the population level, owing to their clustered activity when an animal is near a boundary (Solstad et al., 2008; Lever et al., 2009). Despite the availability of direct intracranial electroencephalography (iEEG) recordings from the human medial temporal lobe during computer-based navigation tasks (e.g., Ekstrom et al., 2005; Watrous et al., 2011; Miller et al., 2013; Vass et al., 2016), no studies thus far have shown direct neural signatures of boundary representation in humans. In the present study, we recorded the local field potential (LFP) from surgical epilepsy patients engaged in a computer-based navigation task, combined with a high-resolution electrode localization method, to investigate boundary-related signals in the human brain in specific subregions of the hippocampal formation (i.e., CA1, Dentate Gyrus, Subiculum, EC, and Perirhinal Cortex). We capitalized on the fact that boundary cells, like other spatial cells, are theta-modulated (e.g., Lever et al., 2009) and that the strength of theta oscillations could indicate neural activity in the hippocampus (McFarland et al., 1975; McNaughton et al., 1983; Rivas et al., 1996; Czurko et al., 1999; Terrazas et al., 2005). We compared oscillatory power in three frequency ranges that have been previously implicated in spatial navigation and memory in humans (Nyhus & Curran, 2010; Watrous et al., 2013; Jacobs, 2014)—1–4 Hz (”low theta” or ”delta”), 4–10 Hz (”theta”), 30-90 Hz (”gamma”)— as subjects encoded targets near or far from the boundaries of the virtual environment.

## Methods

### Participants

The subjects in our study were 58 epilepsy patients between the ages 18 and 65, who had electrodes surgically implanted to localize seizure foci and guide potential surgical treatment. Subjects performed a virtual navigation task on a laptop computer as their neural activity was recorded at 500 Hz or above (Jacobs & Kahana, 2010). Electrodes were implanted in various brain regions as dictated by clinical needs; for the analysis of neural measurements, we selectively analyzed 39 of the patients who had electrodes in our five regions of interest: CA1, Dentate gyrus, subiculum, entorhinal, and perirhinal cortex. These regions were chose because they were the top five hippocampal subregions with the most number of electrodes implanted; 39 subjects had electrodes in those regions. The same methods were applied at seven testing sites: Thomas Jefferson University Hospital (Philadelphia, PA), Mayo Clinic (Rochester, MN), University of Texas Southwestern (Dallas, TX), Geisel School of Medicine at Dartmouth (Hanover, NH), University of Pennsylvania Medical Center (Philadelphia, PA), Emory University Hospital (Atlanta, GA), and Columbia University Medical Center (New York, NY). Each subject provided informed consent prior to participation. Our multi-site study was approved by local institutional review boards (IRBs), as well as the IRB of the University of Pennsylvania (data coordination site) and the Human Research Protection Office (HRPO) at the Space and Naval Warfare Systems Command Systems Center Pacific (SPAWAR/SSC). Data from five subjects who responded randomly in the task (see below for details) were excluded from our analysis.

### Spatial Navigation Task

Subjects performed a computer-based spatial memory task (Jacobs et al., 2016) in a virtual rectangular arena (approximately equivalent to 19 m x 10.5 m) with four distal visual cues for orienting. Each trial (48 trials per session, 1-3 sessions per subject) began with a 2 s period during which subjects were presented with a still scene of the environment. Then, a target object appeared on screen and subjects were automatically rotated (1 s duration) and driven toward (3 s duration, constant speed), until they were stopped at the target location (1 s duration). This 5 second long encoding period took place twice, from two different viewpoints in the environment. Then, after a 5 s delay, subjects were transported to a different location/direction (chosen randomly from a range of locations that would fit the 3 seconds of driving and 1 second of rotating) from which they had to drive themselves back using a joystick to the now hidden target and press a response button. The two encoding periods were separated by a 5 s black screen. Subjects received feedback on their responses by means of a simple rectangular depiction the environment with the target and response locations marked as circular points (see Figure 1a). The automatized design of the encoding phase ensured that all aspects of a subject’s movement (e.g., time, speed, distance, visual flow, movement) were identical across trials (and across target locations), while maximizing the number of trials.

**Figure 1:**
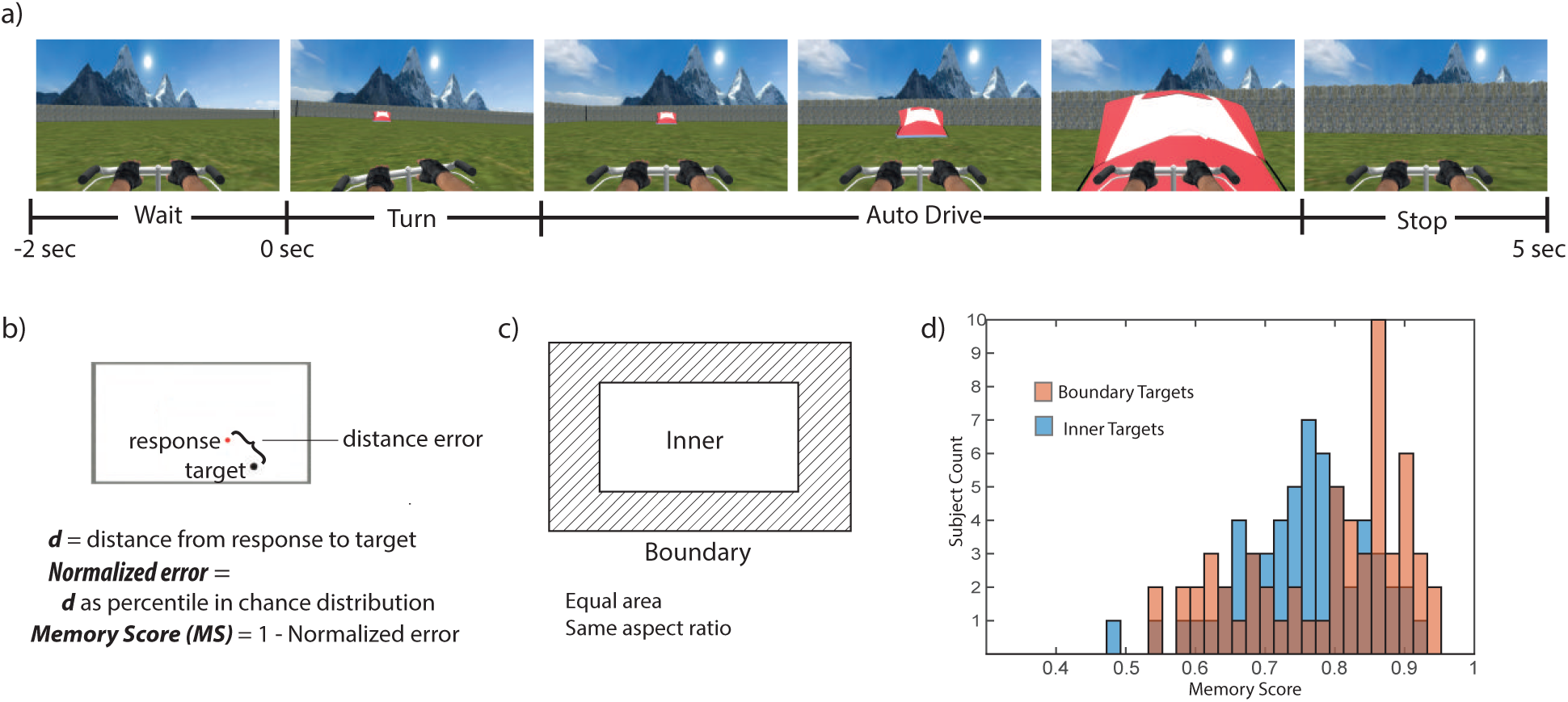
a) Each trial began with a 2 second stationary wait period in which the subject viewed the environment. Once the target object appeared, the subject had 5 seconds of encoding, during which the subject (in the VR environment) was automatically rotated and then driven to the target location. This was repeated from a different starting point, such that there were two encoding trials for each object location. Following the encoding trials, subjects were transported to a new starting point for the test phase and asked to drive themselves back to the goal location and respond by pressing a button. They were shown a map of the goal and their response location and rewarded points accordingly. b) We computed a Memory Score (MS) based on the accuracy percentile, with respect to the chance distribution of responses for each goal location. c) We categorized target locations as being ”Boundary” or ”Inner” by dividing the rectangular environment into two equal areas with equal aspect ratios. d) Subject-wise distributions of memory scores for Boundary and Inner trials, which indicate that subjects performed better on Boundary trials overall.

### Electrode Localization

Prior to surgical electrode implantation, we acquired high-resolution structural magnetic resonance imaging (MRI) scans of the hippocampus and medial temporal lobe from each subject (0.5 mm by 0.5mm by 2mm). The hippocampal subregions and extra-hippocampal cortical regions were automatically defined and labeled on these scanned images using a multi-atlas based segmentation technique (Wang et al., 2013; Yushkevich et al., 2015). After the electrode implantation, a neuroradiologist identified each electrode contact using a post-implant CT scan. The MRI and CT scans were co-registered using Advanced Normalization Tools (Avants et al., 2008), and the neuroradiologist visually confirmed and provided additional detail on the localization and anatomical label for each contact (see Duvernoy, 2005).

### Statistical Analysis of Behavior

We measured patients’ memory performance in a way that accounted for unequal distribution of possible distance errors across the environment. An example of this issue is that objects at the far ends of the environment have a larger maximum possible error distance compared to objects in the center. In our approach we measured performance for each response by computing a memory score (MS), which normalizes for overall difficulty across target locations by computing the actual response’s rank relative to a distribution of a chance distribution based on 100,000 randomly-generated response locations. This means that MS of 1 corresponds to a perfect response (0 error), MS of 0 corresponded to the worst possible response, and MS of 0.5 was chance). We then divided the environment into two equal regions (an outer rectangular ring (”Boundary”) and a central area (”Inner”) and compared subjects’ mean MSs between the two zones (see Figure 1c-d). In order to select only those subjects who understood and were able to perform the task, we discarded data from five subjects who performed overall at chance level (t-test against chance MS of 0.5, n.s.).

### Statistical Analysis of Neural Signals

To compare neural activity between trials in which subjects were successfully encoding both *Boundary* and *Inner* locations (rather than being disoriented or inattentive), we first deleted all trials in which subjects did not successfully encode spatial location (MS < 0.73, chosen based on the mean MS across all subjects) in order to to reduce noise. Furthermore, we only included subjects that had at least 5 trials in each category (Boundary/Inner), in order to ensure sufficient sampling.

We extracted the oscillatory power at each electrode in three frequency bands: low-theta (1–4 Hz), theta (4–10 Hz), and gamma (30–90 Hz). The signals were bandpass filtered in these ranges using a Butterworth filter, notch-filtered at 60 Hz, and the amplitude was extracted with a Hilbert transform (Freeman, 2007). We then computed the time-averaged power in each band across the 5-second encoding period. The power values were then z-scored according to the mean power in that electrode over all encoding periods in the session.

We averaged the power over all electrodes from the same hemisphere of a single patient, such that we took one measurement from each hemisphere and used these as the input to our analysis of variance. This was done in order to prevent double-sampling from shared signal sources within different contacts on a subject’s medial temporal lobe within one hemisphere. Using this method, we had 18 subiculum hemispheric samples (12 in left hemisphere (LH)) from 15 patients (i.e., 3 patients with bilateral subiculum electrodes), 18 entorhinal samples (13 from LH) from 14 patients, and 43 CA1 samples (25 in LH) from 37 patients, 23 dentate gyrus samples (15 in LH) from 21 patients, and 29 perirhinal samples (20 from LH) from 22 subjects.

## Results

### Behavior

If boundaries are crucial to the neural representation of spatial location, subjects should generally be more accurate in their performance for locations near boundaries than for locations far from boundaries (Hartley et al., 2004). To test the degree to which subjects rely on the environmental boundaries to perform our task, we divided the environment into two regions with equal areas (an outer rectangular ring (”Boundary”) and a central area (”Inner”), see Figure 1c) and compared subjects’ performance as quantified by their memory scores (MS) (see *Methods* for details). A pairwise t-test revealed that subjects performed better in *Boundary* trials than in *Inner* trials (t(57)=3.74, p=0.0004; see Figure 1d). These results are consistent with the interpretation that boundaries have a significant influence on the computation of spatial location and indicate that our virtual-reality task sufficiently engaged those underlying navigational mechanisms.

### Neural Results

A repeated-measures analysis of variance was conducted to examine at the population level whether neural signals at three frequency ranges (1–4 Hz, 4–10 Hz, and 30–90 Hz; within-subjects measure) varied according to the presence of a nearby boundary (within-subjects measure) across five different subregions of the hippocampus (CA1, Dentate Gyrus, Subiculum, EC, Perirhinal Cortex) (Figure 2a).

**Figure 2:**
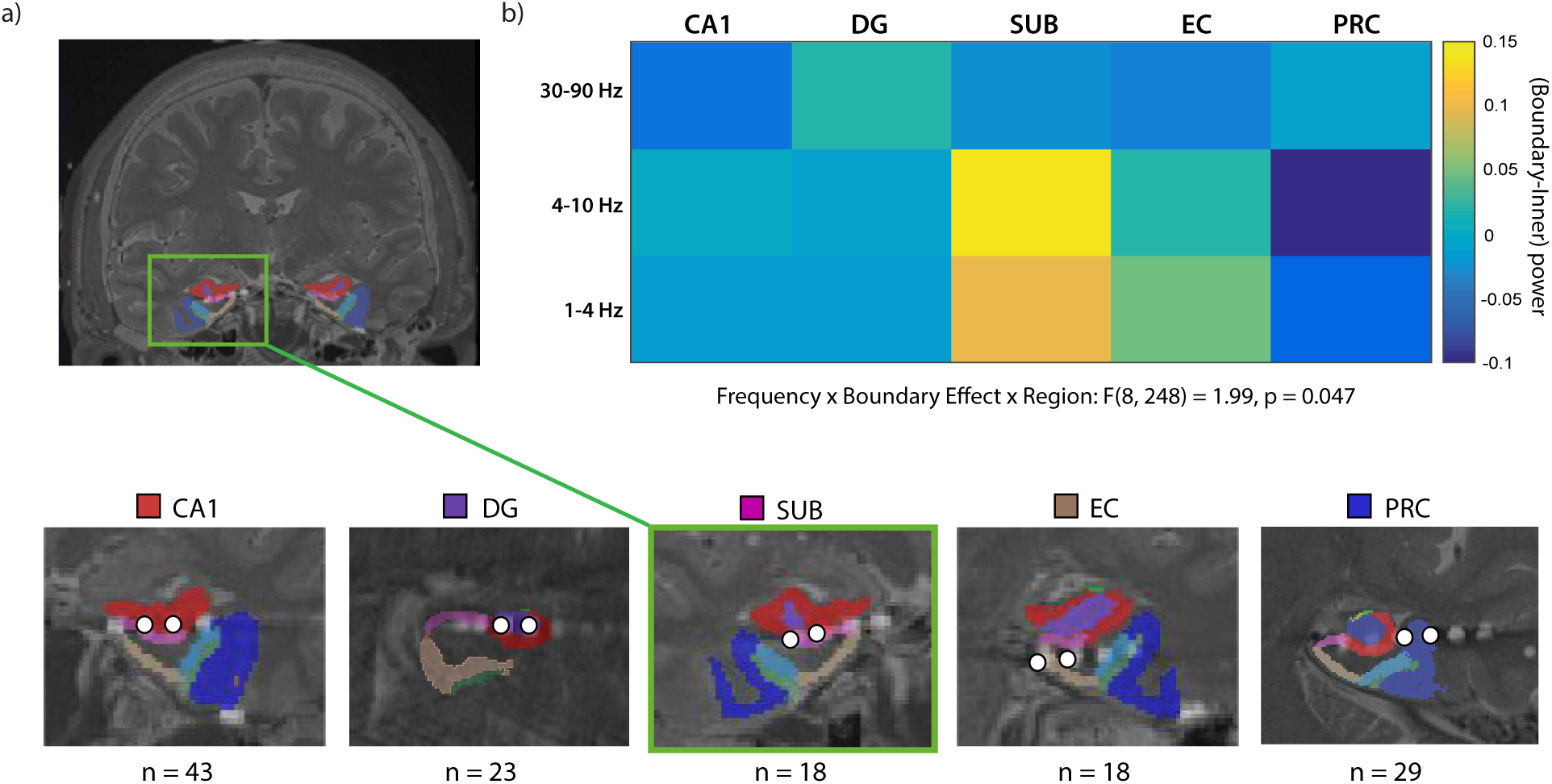
a) Electrodes were localized by combining pre-surgical high-resolution structural MRI and post-implant CT scans. The hippocampal subregions are labeled and shaded color in each example image, and the bipolar electrode contact pairs (distance 1.5 mm) centered at each subregion are marked with white dots. (Images from patient 1066P) b) Encoding near-boundary locations elicited higher power in low frequency oscillations than inner locations. Z-scored power differences between boundary and inner target locations are plotted for three frequency bands (low-theta: 1–4 Hz; theta: 4–10 Hz; gamma: 30–90 Hz). There was a significant Boundary × Frequency interaction that was specific only to the subiculum (F(2,34)=4.71, p=0.016) and present in no other region.

Critically, we found that LFP power across all electrodes significantly varied according to whether the patient encoded a location near or far from a boundary, at particular frequency ranges and localized to a particular region (Boundary × Frequency × Region interaction: F(8,248)=1.99, p=0.047,) (Figure 2b). Upon closer inspection, the Boundary × Frequency effect was specific to the subiculum (F(2,34)=4.71, p=0.016, eta-squared=0.22) and significant in no other region (all F’s < 2, p’s> 0.2). Post-hoc t-tests (Bonferroni corrected) revealed that encoding locations near boundaries elicited greater low-theta and theta power than encoding inner locations (1–4 Hz: t(17)=2.58, p=0.057; 4–10 Hz: t(17)=3.22, p = 0.015; Figure 3a). This effect was not present in the 30–90-Hz gamma band (t(17) < 1, n.s.).

**Figure 3:**
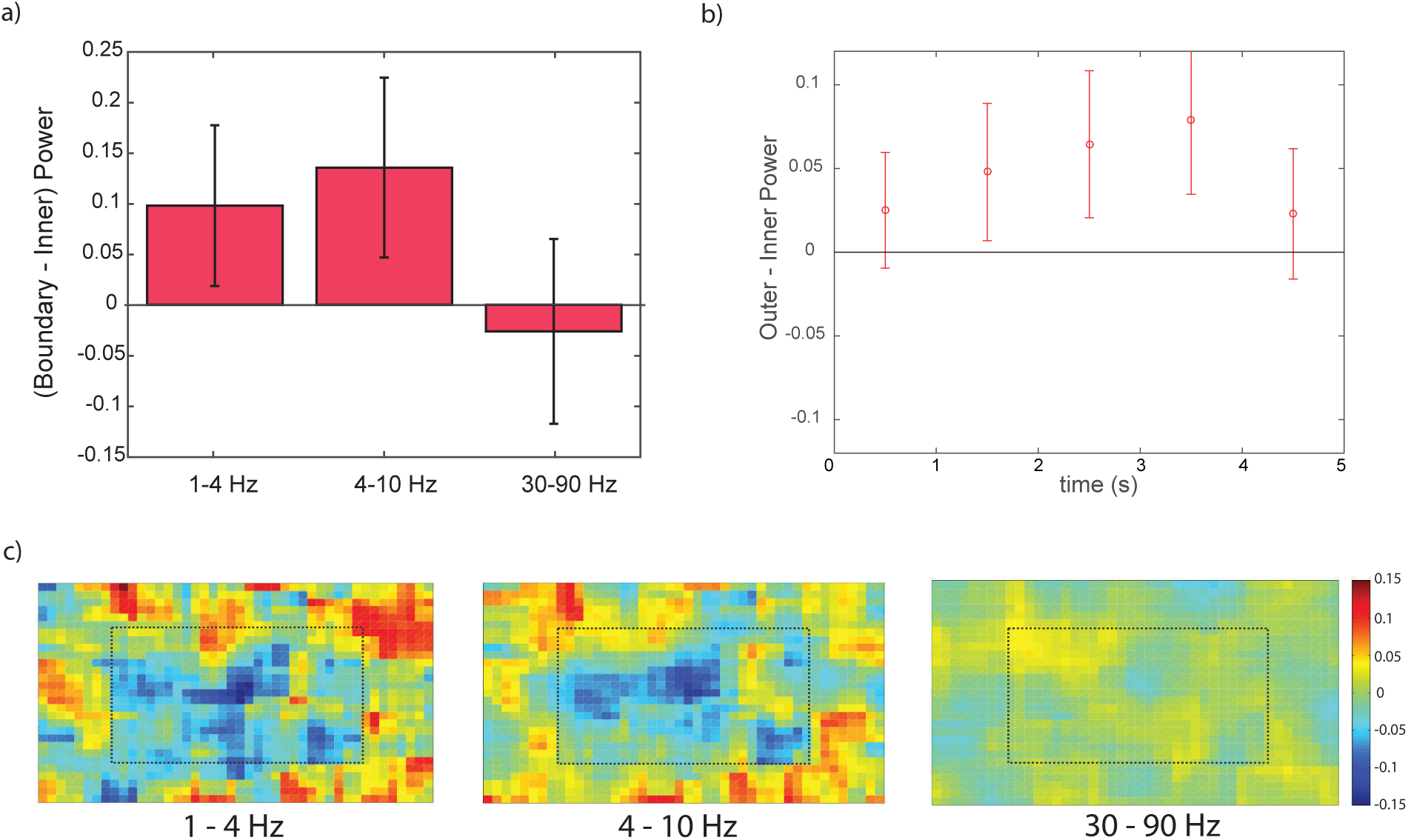
a) Normalized power difference in the subiculum between *Boundary* and *Inner* locations in three frequency bands (low-theta: 1–4 Hz; theta: 4–10 Hz; gamma: 30–90 Hz). Error bars indicate 95% confidence intervals. Goal locations near boundaries elicit stronger theta oscillations than those far from boundaries. (b) Boundary-Inner theta power across the five seconds of navigation to the target. (c) Overhead heat-maps of the environment plotting average z-scored power for the three observed frequency bands. We binned the environment into a 45 by 30 rectangular grid and computed average power in each bin for each subicular sample. Individual heat maps were smoothed with a 2D gaussian kernel (width=7) and then averaged across all samples. Dotted lines indicate the *Boundary-Inner* division.

We next examined this boundary-related signal at the level of individual hemispheric measurements and subjects. 15 out of 18 hemispheric subiculum measurements showed greater theta power for navigating to boundary locations than inner locations (binomial test, p=0.008); this was in 12 out of 15 subjects (binomial test, p=0.035). In some cases, the boundary effect was significant even at the single-session level (see Figure 4). Across all electrodes, there was a significant negative correlation between LFP power on each trial and distance to the closest boundary, for both low theta (1–4 Hz, p = 0.01) and theta (4–10 Hz, p = 0.01), showing that the target location’s proximity to the environmental walls elicited stronger signals in those frequencies (Figure 3b); this indicates that the boundary effect found above is not an artificial consequence our particular designation of *Boundary* and *Inner* regions. Because only 6 out of the 18 subicular samples were from the right hemisphere, we did not have sufficient power to test for hemispheric differences.

**Figure 4:**
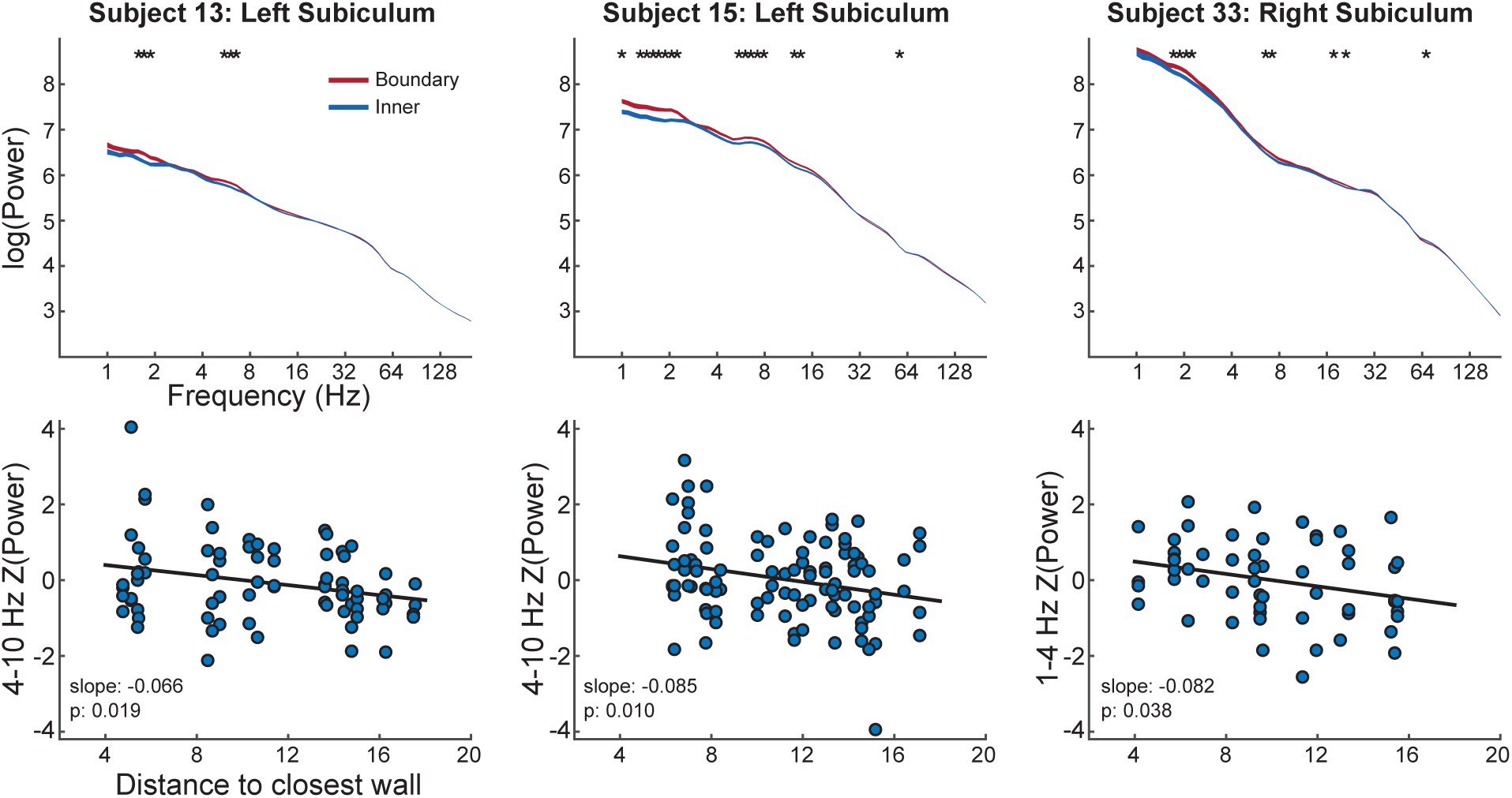
a–c) Each column shows examples of an individual electrode from three different subjects. First row: Examples of power spectra from individual electrodes for *Boundary* and *Inner* trials. Line thickness indicates standard error. Asterisks indicate parts of the spectra where *Boundary* and *Inner* trials significantly differ (t-tests at each frequency, p<0.05). Second row: Trial-by-trial plot of power in theta frequency oscillations for each corresponding electrode above it. Slopes of the best fit lines that negatively deviate from zero show that theta power is stronger at closer distances to a wall boundary.

Although we excluded the trials with memory scores below mean performance (MS< 0.73) in the above analyses, it is still possible that the difference in theta power between the boundary categories simply reflected performance differences between *Boundary* and *Inner* locations (as opposed to boundary proximity per se). If this were true, we should also see theta effects related to performance within the same boundary category (i.e., even for just *Inner* locations, good trials should exhibit higher theta power than bad trials). To test this, we compared LFP power between the ”good” trials (MS> 0.73) and (the previously excluded) ”bad” trials (MS< 0.73), separately for each boundary category. We found no performance effects, for both *Boundary* and *Inner* trials (all t’s≤1.2, n.s.), suggesting that subicular theta was not linked directly to performance. Furthermore, there were no significant boundary-inner differences in the ”bad” trials, suggesting also that the theta effect is not simply driven by perceptual stimulus/visual differences between the boundary and inner trials. Interestingly, however, when comparing across performance categories, even the ”bad” boundary trials elicited significantly higher theta power than the ”good” inner trials, t(17)=2.57, p=0.02 (t< 1.1 for low-theta and gamma).

As a further control to ensure that the difference between boundary and inner trials is truly driven by the target, we analyzed the LFP signals during the first two seconds of each trial, when the subject was standing within the virtual environment and waiting for the target object to appear (−2 to 0 seconds prior to the encoding period; Figure 1a). As expected, before the target object actually appeared on screen, there were no differences between the boundary or in the inner region (all t’s< 1, n.s.). Additionally, to examine whether neural activity varied with respect to the subject’s own location during that two-second period, we compared LFP as a function of the subject’s position (rather than the target position) in the virtual space. We found no effects of boundary proximity in any frequency band (all t’s< 1.5, n.s.). These results are consistent with the interpretation that subjects mainly represented the goal location during the encoding phase of this task.

## Discussion

Our analysis of the local field potential at various hippocampal subregions reveals for the first time in humans that the subiculum may play a key role in boundary-based spatial mapping. This finding extends previous neuroimaging studies implicating the human hippocampus in boundary-based navigation (Doeller et al., 2008; Bird et al., 2010) and is convergent with single-unit recording of boundary cells in the rodent subiculum (Lever et al., 2009). Interestingly, although rodents have boundary-related cells in both subiculum (Lever et al., 2009) and the EC (Solstad et al., 2008), we found boundary-related effects only in the former. One potential explanation for this difference is that the subiculum is more strongly involved than the EC in boundary-based spatial mapping. This account is consistent with rodent studies that reported a much higher percentage of boundary cells in the subiculum: (20–25% Lever et al., 2009; Olson & Nitz, 2017) than EC (6–11% Solstad et al., 2008; Boccara et al., 2010; Bjerknes et al., 2014; Tang et al., 2014).

Could the increase in theta power for boundary-encoding be the result of increased activity from populations of human subicular boundary cells? Unlike other spatial cell types, which activate fairly evenly across an environment, the entire boundary-cell network is more active overall when representing particular areas of an environment (i.e., near boundaries). This coarse-grained spatial specificity in its firing properties is essential for our identification of boundary-related LFP activity. The theta effect we observed might be broadly interpreted as a manifestation of boundary-based spatial encoding or navigation strategies. At the same time, however, it could signify the existence and dynamic activation of boundary-coding cells in the human subiculum. It is noted, however, that this type of boundary-related LFP signal change has not yet been analyzed in rodents, perhaps due to logistical confounds in performing this comparison, such as the variable behaviors of rodents across the environment. For instance, animals run at higher mean speeds parallel to walls (Horev et al., 2006) and theta power increases with running speed (Rivas et al., 1996; Czurko et al., 1999; Maurer et al., 2005), potentially making it difficult to isolate boundary-related theta effects in rodents.

The boundary-related theta patterns we observed appeared during the encoding period of our task when the target location which was visible for 5 s as the subject was automatically moved towards it at a fixed velocity. Although this task design is different from traditional tests of navigation, we implemented this fixed-movement encoding period because it allowed us to equate for multiple perceptual, behavioral, and motoric factors across all trials and for all subjects (Jacobs et al., 2016). In other words, the boundary effect here cannot be attributed to differences in path length or shape, joystick control, speed, visual flow, timing and trial length, just to name a few, between trial with boundary and inner targets. For the same reasons, we have chosen to analyze the encoding phase rather than the freely-moving response phase, in which none of the above factors could be controlled.

The fact that our findings are specific to the target location, rather than the goal location, adds to the body of evidence suggesting a role for the hippocampal formation in attended, viewed, imagined, and planned spatial mapping (Rolls, 1999; Hok et al., 2007; Killian et al., 2012; Pfeiffer & Foster, 2013; Horner et al., 2016; Bellmund et al., 2016) and goal representation (Howard et al., 2014; Chadwick et al., 2015). Under different circumstances, however, it may be possible to detect boundary encoding with respect to self-location rather than the goal location. Our task required subjects to maximally attend to the location of the target, for the five brief seconds that it was on screen; moreover, the automated movement during the encoding period made it unnecessary for subjects to attend to their own navigation through space. A different task design requiring subjects to track and control their own position may detect boundary representations with respect to self location (Ekstrom et al., 2003; Jacobs et al., 2013).

Past studies using human intracranial recordings have demonstrated the involvement of both slow theta (1–4 Hz) and fast theta (4–10 Hz) during both real and virtual navigation (Aghajan et al., 2016; Bohbot et al., 2017); and low frequency oscillations seem to be functionally involved in human memory and navigation (Watrous et al., 2013; Jacobs, 2014). Our boundary-related effects were most strongly seen in the conventional 4–10-Hz range where theta oscillations are commonly found in rodents, but we did also observe a trend, albeit more weakly, at 1–4 Hz. It is possible that both low-theta and theta bands may be implicated in spatial processing in humans. The present results may provide insight that guides future work on identifying potential functional differences between these two bands.

Another open question for further study involves the dissociation of theta power increases in the medial temporal lobe related to memory performance with those related to spatial representation (as we have found in this study). It is difficult to completely disentangle spatial encoding of boundaries to spatial memory, given that successful performance should be a functional consequence of successful spatial encoding, after all. Nevertheless, our control analyses show that the theta increases that we have observed in this task are not solely attributable to memory performance. Moreover, the detailed localization of these effects to the subiculum make it unlikely that these theta effects are actually related to memory (which is a hippocampus-wide phenomenon) rather than to spatial boundaries.

Spatial mapping is one of the most essential survival skills for any self-locomoting animal, and accurate metric representation of distance is essential to accurate place mapping; environmental boundaries, even in naturalistic terrains, provide a stable, invariant cue by which distance representations can be anchored and corrected (Gallistel, 1990). Researchers have suspected for nearly 70 years that even distantly related species like rats and humans share cognitive and neural mechanisms that support such abilities (Tolman, 1948; O’Keefe & Nadel, 1978), and our results fill an important gap in the literature by identifying for the first time a highly-localized neural representation of environmental boundaries in the human subiculum, just as in rats. Not only do these findings inform theories of common spatial coding in the vertebrate brain, they also give us another neural signature which we can use to investigate the flexible application of basic hippocampal representations in supporting abstract human conceptual knowledge (Spelke et al., 2010; Jacobs & Lee, 2016; Constantinescu et al., 2016; Garvert et al., 2017) and the cognitive impairments that result from their dysfunction (Bird et al., 2009; Lakusta et al., 2010).

## Acknowledgments

The authors acknowledge support from the DARPA Restoring Active Memory (RAM) program (Cooperative Agreement N66001-14-2-4032) and NIH Grants MH104606, MH061975. The views, opinions and/or findings expressed are those of the author and should not be interpreted as representing the official views or policies of the Department of Defense or the U.S. Government.

